# Temporal Application of Lysyl Oxidase during Hierarchical Collagen Fiber Formation Differentially Effects Mechanics in Engineered Tissues

**DOI:** 10.1101/2022.10.18.512696

**Authors:** Madison Bates, Leia Troop, M. Ethan Brown, Jennifer L. Puetzer

## Abstract

The primary source of strength in musculoskeletal menisci, tendons, and ligaments are hierarchical collagen fibers; however, these fibers are not regenerated after injury nor in engineered replacements, resulting in limited repair options. Collagen strength is reliant on fiber alignment, density, diameter, and crosslinking. Recently, we developed a culture system which guides cells in high-density collagen gels to develop native-like hierarchically organized collagen fibers, which match native alignment and fiber diameters by 6 weeks. However, tissue mechanics plateau at 1 MPa, suggesting crosslinking is lacking. Collagen crosslinking is regulated by lysyl oxidase (LOX) which forms immature crosslinks that condense into mature trivalent crosslinks. Trivalent crosslinks are thought to be the primarily source of strength in fibers, but its not well understood how they form. The objective of this study was to evaluate the effect of exogenous LOX treatment at different stages of hierarchical fiber formation in our culture system to produce functional engineered replacements and to better understand factors effecting collagen crosslink maturation. We found LOXL2 treatment did not restrict hierarchical fiber formation, with constructs still forming aligned collagen fibrils by 2 weeks, larger fibers by 4 weeks, and early fascicles by 6 weeks. However, LOXL2 treatment did significantly increase mature pyridinium crosslink accumulation and tissue mechanics, with timing of LOXL2 supplementation during fiber formation having a significant effect. Overall, we found one week of LOXL2 supplementation at 4 weeks produced constructs with native fiber organization, increased PYD accumulation, and increased mechanics, ultimately matching the tensile modulus of immature bovine menisci.

**Statement of Significance:** Collagen fibers are the primarily source of strength and function in connective tissues throughout the body, however it remains a challenge to develop these fibers in engineered replacements, greatly reducing treatment options. Here we demonstrate lysyl oxidase like 2 (LOXL2) can be used to significantly improve the mechanics of tissue engineered constructs, but timing of application is important and will most likely depend on degree of collagen organization or maturation. Currently there is limited understanding of how collagen crosslinking is regulated, and this system is a promising platform to further investigate cellular regulation of LOX crosslinking. Understanding the mechanism that regulates LOX production and activity is needed to ultimately regenerate functional repair or replacements for connective tissues throughout the body.

## 1. Introduction

Collagen is a major structure protein which plays an important role in connective tissue biomechanics. In particular, musculoskeletal tissues such as menisci, tendons, and ligaments derive their strength and function primarily from large type I collagen fibers that run the length of the tissue. In the meniscus, these collagen fibers run circumferentially around the tissue, helping to resist hoop stress and providing the strength necessary to distribute compressive loads across the knee [1,2]. In tendons and ligaments these collagen fibers are the predominate component of the tissue (~70-80% by dry weight), providing the strength necessary to translate loads from muscle-to-bone and bone-to-bone, respectively [3,4]. Cells in these tissues organize the collagen hierarchically, assembling tropocollagen molecules into nanometer wide fibrils, micron wide fibers, and millimeter to centimeter wide fascicles, growing in size and strength with increasing mechanical demand [1,3,5–11].

When these musculoskeletal tissues are injured, the collagen fibers are torn resulting in loss of function, pain, decreased mobility, and lost work days [2,4,12,13]. These tissues are characterized by little to no healing, ultimately resulting in over 24 million doctor visits for sprains or strains each year and 1.4 million surgeries a year in the US (Based on 1,000,000 meniscus, 135,000 ACL, and 275,000 rotator cuff repairs per year) [2,4,12–14]. Currently, small tears are treated by various rehabilitation protocols, which often yield scar tissue with unorganized collagen and inferior mechanics [15–17]. More severe tears are replaced by autograft or allograft transplants, which have limited availability, risk of immune response, donor site morbidity, and high re-rupture rates [2,4,12,14,18]. Ultimately, current treatments for torn meniscus, tendon, or ligament, whether repaired or replaced, fail to restore the structural and functional properties of native tissue [2,4,12,15,18]. Engineered tissues are a promising alternative for repairing these musculoskeletal tissues [2,4,12], but it remains a challenge to regenerate the organized collagen fibers that are essential to attaining native mechanics. Thus, often these engineered replacements do not translate to the clinic. It is poorly understood how these fibers develop *in vitro*, and knowing more about this process could help to further develop orthopaedic engineered tissues as a whole.

Recently, we developed a novel culture system which guides meniscus, tendon, and ligament cells in high density collagen gels to produce native-like hierarchically organized collagen fibers (**Figure 1B**) [19–22]. Specifically, compressive boundary restraints (clamps) applied at the edge of the gel restrict cellular contraction and guide cells to form aligned collagen fibrils by 2 weeks, native-sized fibers by 4 weeks, and 50-350 μm wide fiber bundles or early fascicles by 6 weeks [19]. The fiber maturation in this system resembles native fibrillogenesis, where collagen fibrils first align and grow longitudinally, then expand laterally [3,7,19]. This culture system yields some of the largest, most organized fibers produced to date; however, tensile properties plateau at ~1 MPa [19] indicating these constructs need further maturation to serve as functional replacements. It has been reported that tissue strength, in native and engineered tissues, is reliant on collagen fiber density, diameter, orientation, and degree of crosslinking [5,23–26]. With our culture system we are able to match native tissue alignment by 2 weeks and native fiber sizes by 4-6 weeks [19]; however, tissue mechanics are still inferior to native tissue, suggesting a need for increased collagen crosslinking.

**Figure 1:**
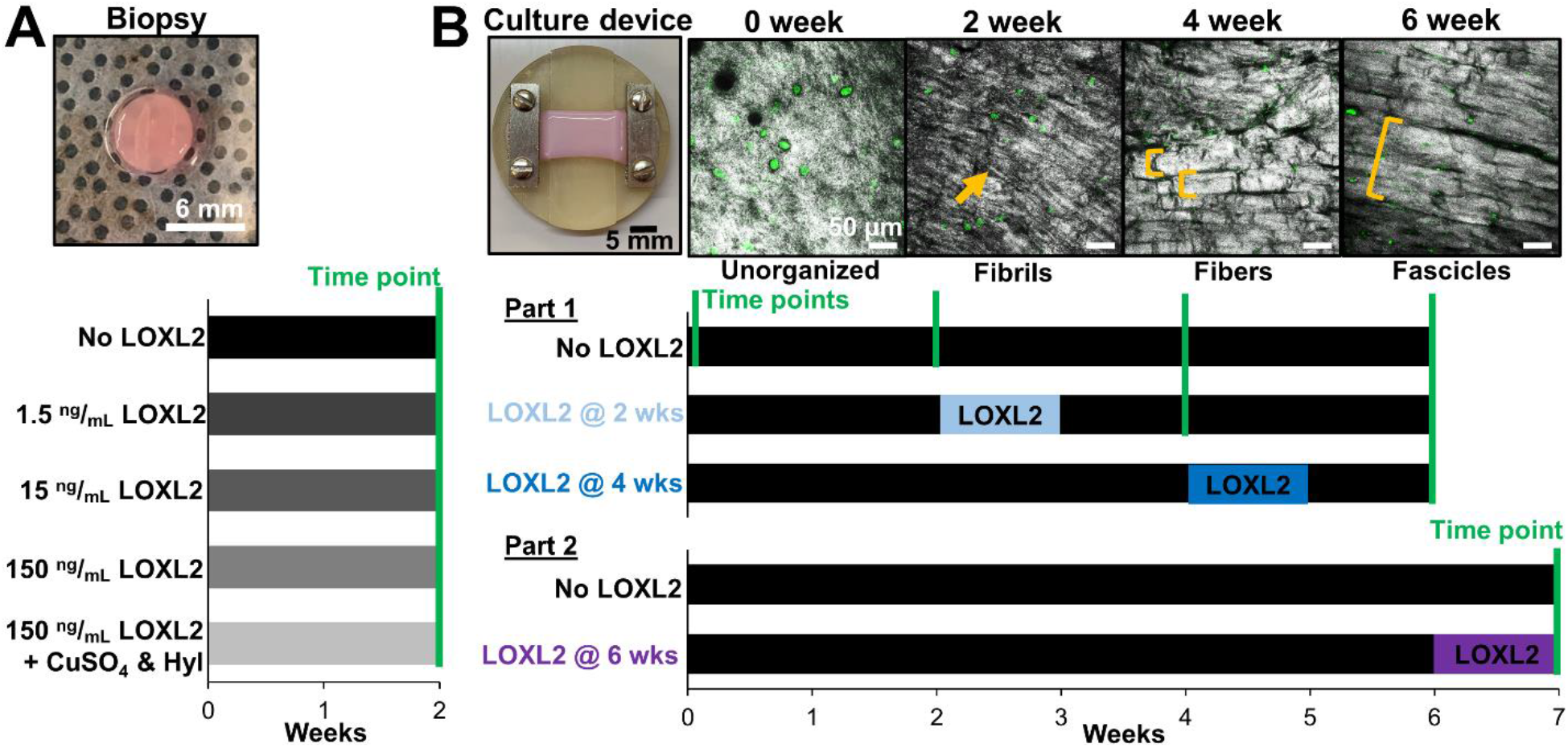
Experimental setup for analyzing the effect of lysyl oxidase like 2 (LOXL2) concentration and temporal application. A) To evaluate the effect of LOXL2 concentration, cell-seeded 6 mm biopsy punches were cultured with 0, 1.5, 15, or 150 ng/ml of LOXL2 for 2 weeks. B) To evaluate the effect of LOXL2 at various levels of collagen organization cell-seeded constructs were cultured in clamping culture device to guide the formation of hierarchical fibers over 6 weeks. Images are confocal reflectance of collagen organization (grey = collagen, green = cellular auto-fluorescence, arrows denote aligned fibrils, brackets denote fibers and fascicles). In Part 1 of the experiment 150 ng/mL LOXL2 was added to the media for one week once constructs developed aligned fibrils (at 2 weeks) and fibers (at 4 weeks) and cultured out to 6 weeks. In Part 2 LOXL2 was added to the media at 6 weeks for one week to evaluate the effect of LOXL2 once fascicles start to form.

Collagen crosslinking is the final step in the synthesis of collagen fibers, and is known to occur during or after self-assembly of fibrils [27–29]. It is mediated by the enzymes lysyl oxidase (LOX) and lysyl oxidase like isoforms (LOXL) which catalyzes lysine and hydroxylysine residues into reactive aldehydes that then produce divalent crosslinks with aldehydes on adjacent molecules. These divalent crosslinks stabilize newly formed fibrils and can eventually condense to form mature trivalent crosslinks by reacting with a third residue or another divalent crosslink [28,30–33]. Trivalent crosslinks increase throughout development as hierarchical collagen fibers mature and are primarily found in highly loaded tissues [28,34,35]. They are thought to be a primary source of strength in collagen fibers [26,28,31,34,36]; however, it is not well understood how they form [28,31,37]. The most common type of trivalent crosslink found in musculoskeletal tissues are Pyridinoline (PYD) crosslinks which encompass both hydroxylysyl pyridinoline (HP, composed of 3 hydroxylysine residues) and lysyl pyridinoline (LP, composed of 2 hydroxylysine residues and 1 lysine) crosslinks [28,33,38]. It has been reported that collagen needs to be pre-assembled into a quarter staggered fibrillar array with the proper aldehyde residues present prior to LOX activation for crosslinks to form [24,28,30,31], demonstrating the importance of collagen organization in enzymatic crosslinking.

Recent studies have shown that scaffold free constructs treated with exogenous lysyl oxidase homolog 2 (LOXL2) have increased trivalent crosslinks and tissue strength [24,39—41]. Therefore, we were interested in investigating whether exogenous LOXL2 treatment would increase mechanical properties in our culture system. Also, based on the importance of fibril alignment and organization to crosslink formation, we were interested in exploring how the underlying collagen organization affects the development of mature trivalent crosslinking in our system. Therefore, the objective of this study was to find an optimal exogenous LOXL2 concentration for inducing crosslinking in high density collagen gels and apply it temporally to our culture system to investigate how degree of hierarchical organization affects LOXL2 induced crosslinking and tissue mechanics. We hypothesize temporal LOXL2 treatment in our culture system as collagen fibrils align and develop into fibers will significantly increase trivalent crosslink accumulation and tissue tensile strength, producing tissues with native levels of collagen organization and strength.

## 2. Materials and Methods

### 2.1 Cell isolation and construct fabrication

Bovine meniscal fibrochondrocytes were used as a generic primary cell source since we have previously demonstrated these cells produce native-sized hierarchical collagen fibers in high density collagen gels, similar to bovine ligament fibroblasts and tenocytes [19]. To obtain meniscal fibrochondrocytes, calf legs (1-6 weeks old) were purchased from a local abattoir within 48 hours of slaughter and the menisci were aseptically isolated from the joint, diced, and digested overnight with 0.3% w/v collagenase (Worthington) as previously described [19,21,22,42,43]. Fibrochondrocytes were isolated from the entire meniscus and cells from 3 bovines were pooled per isolation to limit donor variability, with at least 3 separate isolations used to create constructs (9 bovines total) [19,42]. Meniscal fibrochondrocytes from each isolation were seeded at 2800 cells/cm^2^ and passaged 1-2 times to yield sufficient cell numbers. Cells were expanded in basic growth media consisting of Dulbecco’s Modified Eagle Media (DMEM), 10% fetal bovine serum (FBS), 1% antibiotic/antimycotic, 0.1 mM non-essential amino acids, 50 μg/mL ascorbic acid, and 0.8 mM L-proline, changed every 3 days [19,21,22,42,43].

Cell-seeded constructs composed of high-density collagen were fabricated as previously described [19–22,42,43]. Briefly, type I collagen was extracted from purchased adult mixed gender Sprague-Dawley rat tail tendons (BIOIVT) and reconstituted at 30 mg/ml in 0.1% v/v acetic acid. The stock 30 mg/ml collagen solution was mixed with appropriate volumes of 1N NaOH and phosphate-buffered saline (PBS) to initiate gelation and return collagen to pH 7 and 300 mOsm osmolarity [21,43,44]. The collagen solution was then immediately mixed with a cell/media suspension, injected between glass sheets 1.5 mm apart, and gelled for 1 hour at 37°C to obtain a sheet gel at 20 mg/ml collagen and 5 x 10^6^ cells/ml. Biopsy punches (6 mm diameter) or rectangular constructs (30 x 8 mm) were cut from the sheet gel and split evenly across time points and experimental conditions. Each sheet gel was made from a single expansion of cells and was created using unique batches of collagen, with each sheet gel yielding 4-6 rectangular constructs.

### 2.2 Experimental setup

Separate experiments were performed to evaluate the effect of exogenous LOXL2 concentration and temporal LOXL2 application during hierarchical fiber development (**Figure 1**). To evaluate the effect of LOXL2 concentration, 6 mm biopsy punches were cut from sheet gels and cultured for up to 2 weeks in the same basic growth media formulation used for cell expansion. Media was supplemented with 0, 1.5, 15, or 150 ng/ml recombinant human LOX homolog like 2 (LOXL2, R&D Systems), with media changed every 2-3 days, similar to previous work in scaffold free and synthetic scaffolds [24,41,45,46]. Since LOX mediated crosslinking is copper dependent, additional constructs were cultured with 150 ng/ml LOXL2, 146 μg/ml hydroxylysine (Hyl), and 1.6 μg/ml copper sulfate (CuSO4) [23,24,39,41,45] to investigate whether copper supplementation is needed in the presence of FBS **(Figure 1A**).

To investigate the effect of exogenous LOXL2 at various levels of hierarchical collagen fiber organization, rectangular constructs were clamped into our culture device on day 1 and cultured for up to 6 weeks to guide cellular hierarchical fiber formation (**Figure 1B**) [19]. Specifically, boundary constraints at the edge of the gel restrict cellular contraction and guide cells to develop aligned collagen fibrils by 2 weeks, native-sized fibers by 4 weeks, and larger collagen bundles or fascicles by 6 weeks, as previously reported [19]. Constructs were maintained in the same basic growth media used for cell expansion and 150 ng/ml recombinant LOXL2 was added to media for 1 week at 2 or 4 weeks of culture, when cells form aligned fibrils and fibers, respectively (**Figure 1B part 1**). Media was changed every 2-3 days, with LOXL2 supplementation occurring three times throughout the week. After one week of LOXL2 supplementation constructs were maintained in basic growth media for up to 6 weeks. Control constructs were cultured without any addition of LOXL2, with time points at 0, 2, 4 and 6 weeks of culture.

Based on the interesting results found with LOXL2 supplementation at the fibril and fiber level, an additional study was performed to evaluate the effect of LOXL2 supplementation once larger collagen bundles or fascicles form. Again rectangular constructs were clamped into our culture device on day 1 and cultured for up to 7 weeks to guide cellular hierarchical fiber formation. At 6 weeks, treated constructs had 150 ng/ml LOXL2 added to the media for 1 week, while controls remained in basic growth media (**Figure 1B part 2**). Time points were taken at 7 weeks of culture.

### 2.3 Postculture analysis

At each time point, 4-8 constructs per group were removed from culture, photographed, weighed, and sectioned into pieces for analysis of collagen organization, composition, and mechanical properties as previously described [19–22,42]. Zero week constructs were harvested on day 2 to allow for 24 hour of clamped culture. The final weights of constructs were normalized to the average construct weight at 0 weeks to determine change in mass with time in culture. N noted in figures represents number of individual constructs analyzed, with each construct being obtained from different sheet gels composed of unique batches of collagen and/or different sets of expanded cells. A variable number of constructs (N = 4-8) were evaluated at each time point due to a large number of analysis techniques and significant contraction with time in culture.

#### 2.3.1 Collagen organization analysis

To evaluate collagen organization, half-sections of biopsy punches or full-length sections of rectangular constructs were fixed in 10% formalin, stored in 70% ethanol, and image with confocal reflectance (N = 4-8). Engineered tissues were compared to native immature (1-6 week old) bovine menisci. Confocal reflectance was performed as previously described [19–22,43]. Briefly, confocal reflectance was performed in combination with fluorescent imaging on a Zeiss 710 Laser Scanning Microscope, using a 405 nm and 488 nm laser, respectively. Collagen organization was assessed by collecting reflected light at 400-469 nm through a 29 μm pinhole with a pixel dwell time of 0.79 μs and cells were imaged by collecting auto-fluorescence at 509–577 nm. Biopsy punches were assessed with a Plan-Apochromat 63x/1.4 Oil DIC M27 objective and rectangular constructs were assessed with a LD C-Apochromat 40x/1.1 W Korr M27 objective. Representative images were taken across the entire full-length section of rectangular constructs at all time points to assess collagen organization across the entire construct.

Following confocal reflectance, native immature menisci and a subset of 6 and 7 week rectangular constructs were analyzed with polarized picrosirius red imaging (N = 3) and scanning electron microscopy (SEM, N = 3) to evaluate fascicle (>100 μm length-scale) and fibril level organization (<1 μm length-scale), respectively. For histological analysis, fixed sections of constructs were embedded into paraffin, sectioned, and stained with picrosirius red. Images were taken on a Nikon Ts2R inverted microscope with a Plan Fluor 10x/0.30 OFN25 Ph1 DLL objective under linear polarized light to assess collagen organization at the fascicle (>100 μm) length-scale as previously described [19,20].

For SEM analysis, fixed sections of constructs were critical point dried, coated with platinum ~25–35 angstroms thick, and imaged with a Hitachi SU-70 FE-SEM at a working distance of 15 mm, 5kV, and 50,000x magnification, as previously described [20,42]. Five representative images from completely separate regions of the construct were obtained from N = 3 separate constructs per condition. Diameters of 15 fibrils per image (total 75 fibrils per construct) were measured with FIJI (NIH) and pooled to determine the average fibril diameter for each construct. Replicate constructs from each group were averaged to determine average fibril diameters and standard deviation between constructs for each treatment group.

#### 2.3.2 Composition Analysis

Biochemical analysis of DNA, glycosaminoglycans (GAG), and collagen content were performed as previously described [19–22,42,43]. Briefly, sections of biopsy punches and rectangular constructs at each time point were weighed wet (WW), frozen, lyophilized, weighed dry (DW) and digested in 1.25 mg/ml papain solution for 16 hours at 60°C (N = 4–8). DNA, GAG, and collagen content were assessed via Quantifluor dsDNA assay kit (Promega), a modified 1,9-dimethylmethylene blue (DMMB) assay at pH 1.5 [47], and a modified hydroxyproline (hypro) assay [42,48], respectively. To account for differences in contraction between time points, biochemical properties were normalized to WW of the samples and multiplied by the total wet weight of the constructs when removed from culture to determine DNA, GAG, and hydroxyproline per constructs.

The same samples used for assessing DNA, GAG, and collagen, were also used to asses accumulation of LOX-mediated trivalent pyridinium (PYD) crosslinks using the MicroVue PYD EIA (Quidel, San Diego, CA) [49]. Briefly, papain digested samples were hydrolyzed in 5N HCL at 110°C for 24 hours, frozen, lyophilized, and reconstituted in deionized water. The MicroVue ELISA was then performed according to manufacture protocol and PYD concentration was normalized to sample wet weight and hydroxyproline content.

As a means to evaluate endogenous LOX produced by cells throughout culture, media from control constructs (N = 4) was collected at each media change every 2–3 days for up to 7 weeks. Media was frozen, stored at-80°C, and analyzed with a fluorometric LOX activity assay (Abcam, ab112139). The assay was performed according to manufactures protocol with media samples taken throughout culture compared to fresh media to determine fold change in active LOX released to the media. Recombinant LOXL2 (R&D Systems) was used as a positive control.

#### 2.3.3 Mechanical analysis

Tensile tests were performed on clamped constructs to assess changes in tissue mechanics with time in culture and temporal LOXL2 application. All mechanical tests were performed using an ElectroForce 3200 System (Bose) outfitted with a 250g load cell, as previously described [19,21,22]. Briefly, full length strips from rectangular constructs were frozen for storage, thawed in PBS with EDTA-free protease inhibitors, measured, and secured in clamps. Samples were loaded to failure at a strain rate of 0.75% strain/s, assuming quasi-static load and ensuring failure between the grips. Tensile properties were determined via a custom least squares linear regression based MATLAB code. The toe and elastic region moduli were determined by fitting the stress-strain curve with a linear regression, ensuring an r^2^ > 0.999. The transition stress and strain were defined as the point where the linear regression of the toe and elastic region intersected. The ultimate tensile strength (UTS) and strain at failure are the maximum stress point.

### 2.4 Statistics

SPSS was used to test for normality of data within each group. After confirming normality, all data were analyzed via 1 or 2-way ANOVA with Tukey’s t-test for post hoc analysis (SigmaPlot 14). For all tests, *p* < 0.05 was considered the threshold for statistical significance. All data are expressed as mean ± standard deviation. Prior to LOXL2 supplementation, all constructs are the same as control non-treated constructs, thus redundant data for LOXL2 treated constructs prior to LOXL2 supplementation has been removed from graphs to aid with visualization of data. However, this data was included in statistical analysis to allow for 2-way ANOVA analysis with time in culture.

## 3. Results

### 3.1 Evaluating optimal LOXL2 concentration in high density collagen gels

We first evaluated the effect of LOXL2 at different concentrations on cell-seeded high-density gels over 2 weeks of culture to determine an optimal concentration for inducing crosslinking while maintain cell viability. Confocal reflectance of constructs after 2 weeks of culture demonstrated that control constructs without LOXL2 treatment maintained unorganized collagen with no distinct collagen fibril formation at the length-scale of analysis. However, in constructs treated with LOXL2 there was increased fibril formation and length of fibrils with increasing LOXL2 concentration (**Figure 2A**), suggesting increased crosslinking.

**Figure 2:**
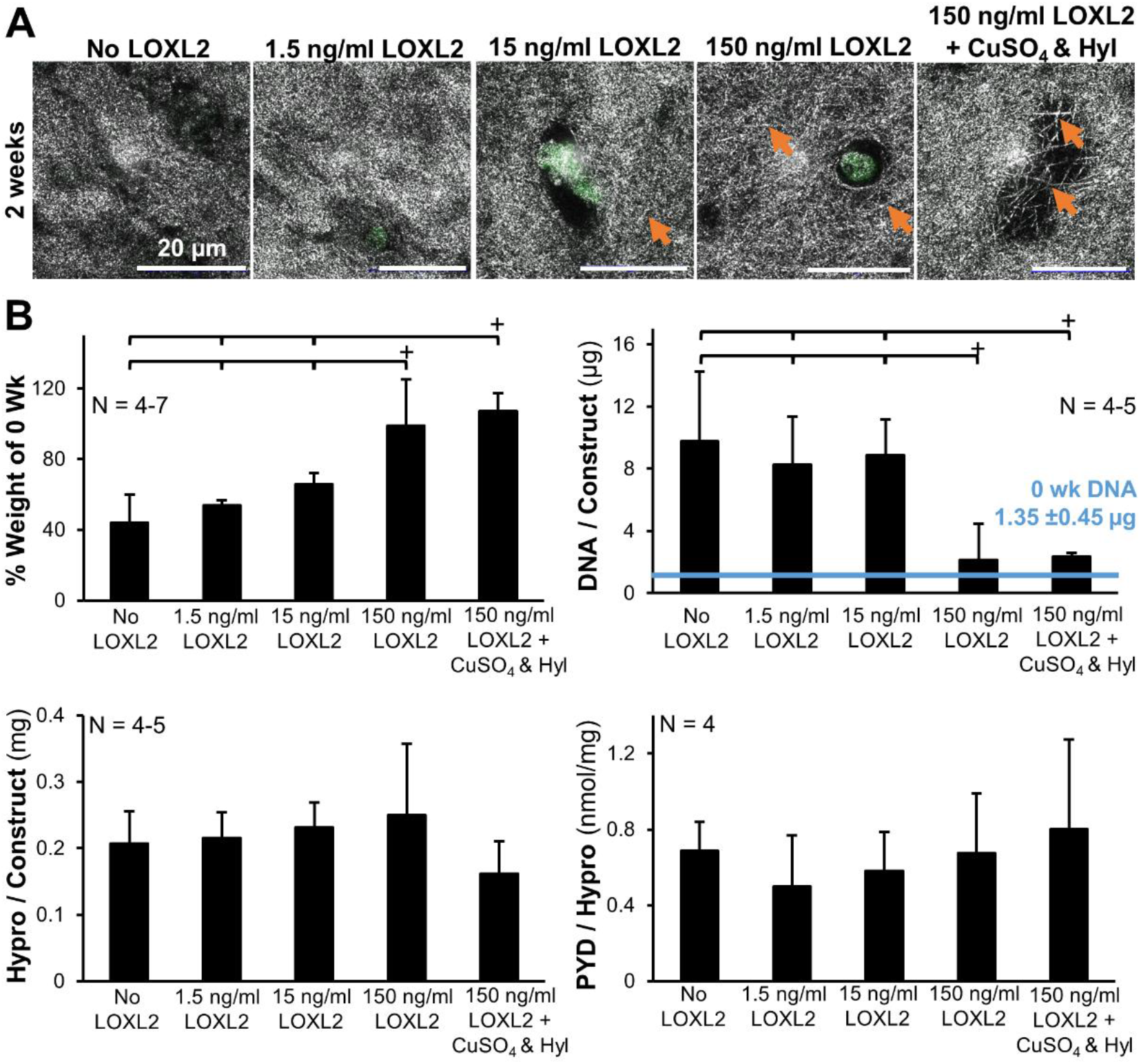
Increasing concentrations of LOXL2 appears to produce dose-dependent increases in divalent crosslinking but does not produce increased trivalent crosslinking in unorganized collagen after 2 weeks of supplementation. A) Confocal reflectance at 2 weeks showing increased fibril formation with increasing LOXL2 concentrations, suggesting increased crosslinking. Grey = collagen, green = cellular autofluorescence, arrows denote fibrils. B) Biopsy punch composition at 2 weeks when cultured with increasing concentrations of LOXL2, including percent weight of constructs compared to weights at 0 weeks, DNA and hydroxyproline (hypro) per construct, and pyridinium (PYD) trivalent crosslinks normalized to hydroxyproline. +significant difference compared to bracketed groups (p<0.05).

Further, while control constructs and constructs treated with low concentrations of LOXL2 had significant decreases in percent weight by 2 weeks of culture, 150 ng/ml LOXL2 treated constructs maintained their size and DNA through 2 weeks of culture, suggesting cellular contraction may be impeded by crosslinks (**Figure 2B**). Of note, control constructs and constructs treated with lower concentrations of LOXL2 did have significantly higher amounts of DNA by 2 weeks of culture, which may suggest higher concentrations of LOXL2 may reduce cellular proliferation. Interestingly, there were no significant differences in mature PYD crosslink accumulation for all LOXL2 concentrations (**Figure 2B**), suggesting exogenous LOXL2 does not produce trivalent crosslinks in unorganized collagen gels over 2 weeks of culture. However, 150 ng/ml LOXL2 did induce fibril formation implying immature divalent crosslinking. Finally, no differences were observed with Hyl and CuSO4 added to the media, suggesting FBS has enough copper to activate LOXL2 [50]. Thus 150 ng/ml LOXL2 was chosen for temporal analysis.

### 3.2 Construct appearance and shape fidelity with temporal LOXL2 application

Gross inspection revealed that all constructs had a reduction in size to ~50% of the original weight by 2 weeks of culture and maintained this size and shape through 6 weeks of culture (**Figure 3**). The addition of LOXL2 at 2 and 4 weeks of culture had no significant effect on gross morphology or size of constructs.

**Figure 3:**
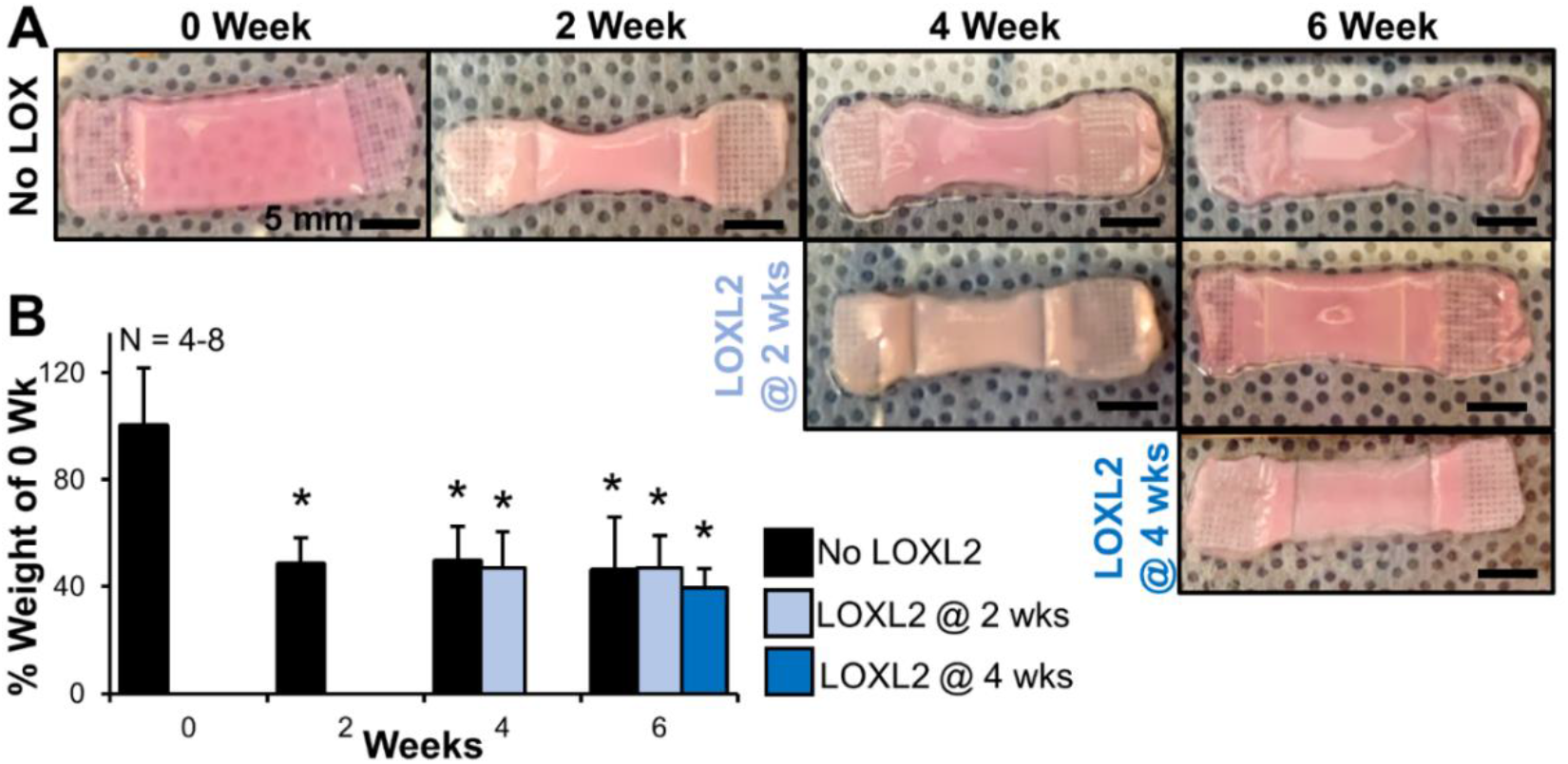
The addition of LOXL2 at 2 and 4 weeks has little to no effect on construct gross morphology and percent weight. A) Gross morphology and B) percent weight of constructs in comparison to construct weights at 0 weeks. *Significant difference compared to 0 week constructs (p<0.05). Prior to LOXL2 treatment at 2 and 4 weeks constructs match control (no LOXL2) constructs, thus redundant data prior to LOXL2 supplementation has been removed to allow for better visualization of the data.

### 3.3 Hierarchical collagen organization with temporal LOXL2 application

Similar to previous studies [19,20], culturing constructs with boundary conditions guided meniscal fibrochondrocytes in high density collagen gels to produce hierarchically organized collagen fibers over 6 weeks of culture (**Figure 4**). Confocal reflectance analysis revealed that all constructs developed aligned fibrils by 2 weeks, which developed into larger collagen fibers ~20- 30 μm in diameter by 4 weeks, and early fascicle-like organizations by 6 weeks, similar to immature bovine menisci (**Figure 4A**). The addition of LOXL2 at 2 and 4 weeks does not significantly alter this collagen organization, however LOXL2 treatment did appear to result in slightly more distinct fiber formation at 4 and 6 weeks, respectively. Similarly, polarized picrosirius red analysis, performed at lower magnification (double length scale), revealed aligned collagen organization in all treatment groups at 6 weeks, with constructs treated with LOXL2 at 4 weeks appearing to have slightly larger, more densely packed collagen fibers than other treatment groups (**Figure 4B**).

**Figure 4:**
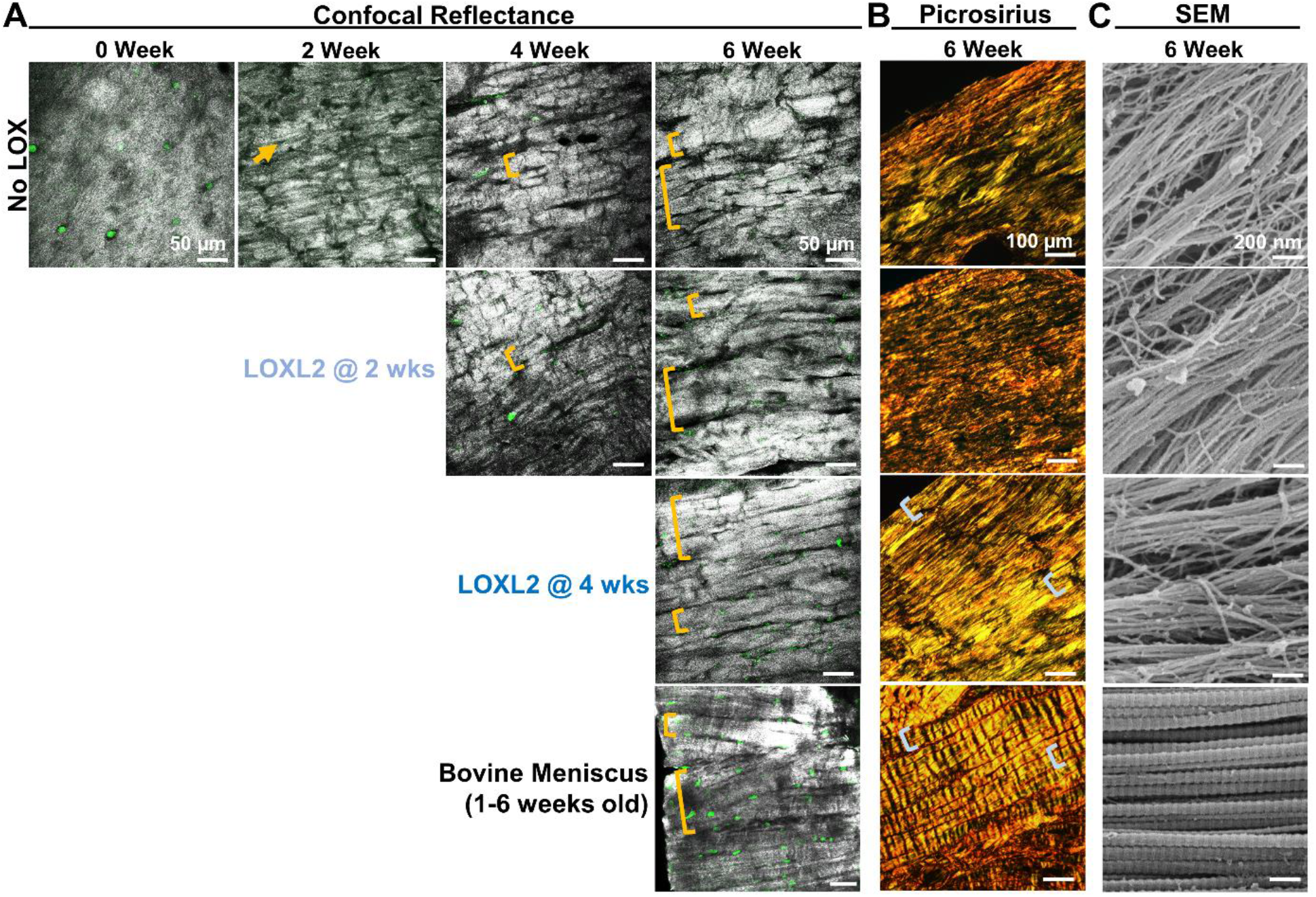
Addition of LOXL2 at 2 and 4 weeks had little effect on collagen organization at the fibril, fiber, and fascicle length-scale. A) Confocal reflectance analysis of fiber-level organization throughout culture (Scale bar = 50 μm, grey = collagen, green = cellular auto-fluorescence, arrows denote aligned fibrils, brackets denote fibers and fascicles), B) Picrosirius red stained sections imaged with polarized light to evaluate fascicle-level organization at 6 weeks (Scale bar = 100 μm) C) SEM analysis of fibril-level organization at 6 weeks (Scale bar = 200 nm).

Fibril level analysis with SEM demonstrated that all treatment groups developed aligned collagen fibrils ~20 nm in diameter which appear to be grouping together into larger bundles, similar in dimensions to that of native meniscus fibrils (**Figure 4C**, immature meniscus fibril diameter 63.8 ±8.3 nm, based on 130 pooled fibril measurements). Interestingly, constructs treated with LOXL2 at 2 and 4 weeks had a significant 5% decrease in fibril diameter compared to constructs not treated with LOXL2, suggesting crosslinking is pulling molecules more closely together and reducing the overall diameter of the fibril (**Table 1**).

**Table 1:** Average fibril diameter of cell-seeded constructs at 6 and 7 weeks

|  | Fibril Diameter (nm) |  |
| --- | --- | --- |
|  | 6 weeks | 7 weeks |
| No LOXL2 | 22.4 ± 0.5 | 21.6 ± 0.1 |
| LOXL2 at 2 weeks | 21.1 ± 0.4<sup>a</sup> | - |
| LOXL2 at 4 weeks | 21.3 ± 0.1<sup>a</sup> | - |
| LOXL2 at 6 weeks | - | 20.9 ± 0.5<sup>a</sup> |

Values are Mean ± SD. Fibril diameter was determined by averaging 75 fibril diameters per construct (15 fibrils measured per SEM image, 5 images per construct), with N = 3 constructs analyzed per time point.
<sup>a</sup> Significance compared to No LOXL2 controls at respective time point ( $p < 0.05$ ).

### 3.4 Tissue composition and crosslink formation with temporal LOXL2 application

Tissue level analysis revealed that DNA and collagen content (represented by hydroxyproline) were largely unchanged with time in culture and with LOXL2 treatment (**Supplemental Figure 1A and Figure 5**). Similarly, GAG accumulation remained relatively steady throughout culture, however constructs treated with LOXL2 at 2 weeks did have significantly reduced GAG accumulation by 6 weeks compared to control constructs (**Supplemental Figure 1B**).

**Figure 5:**
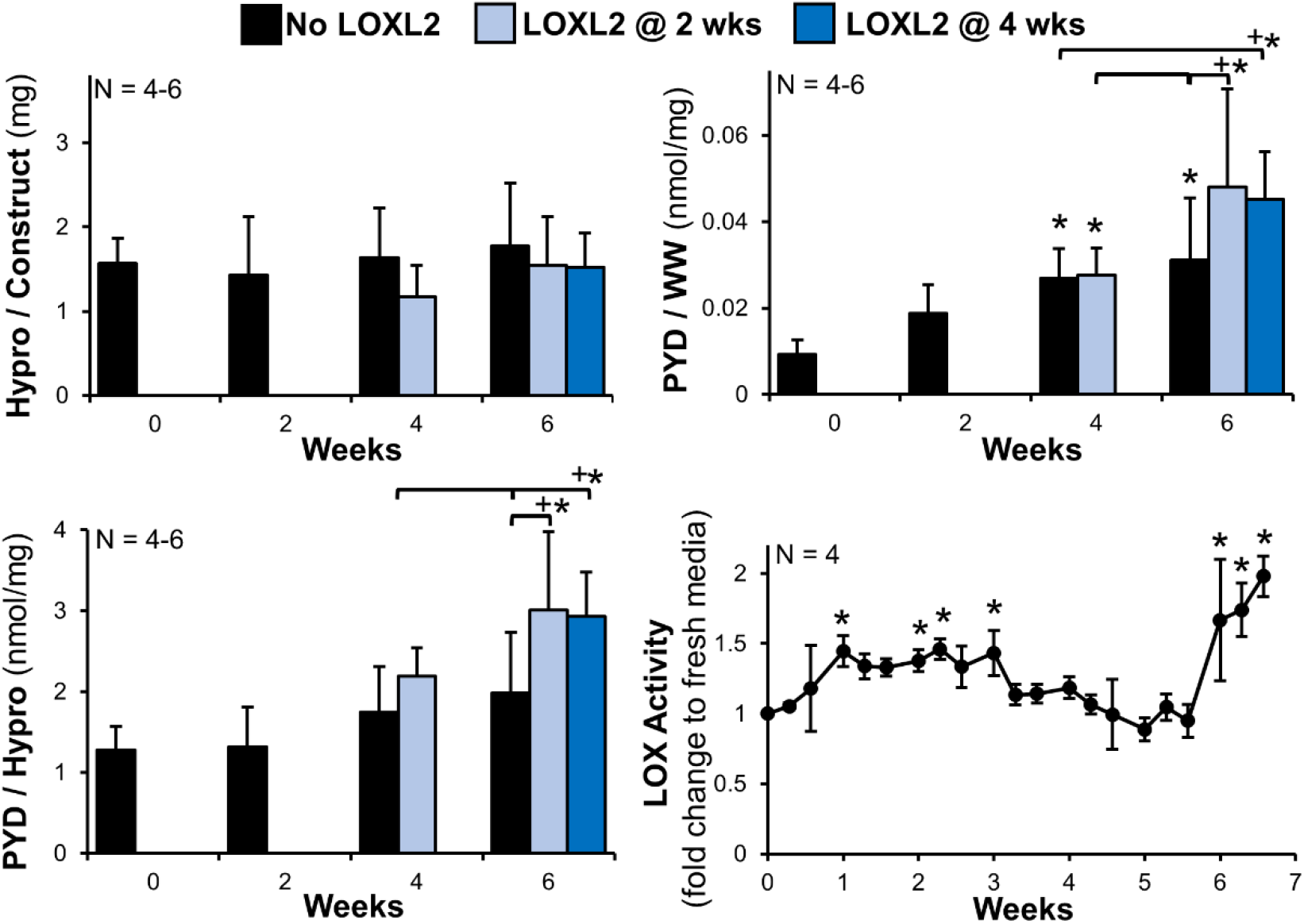
LOXL2 at 2 & 4 weeks of culture has a significant effect on accumulation of trivalent pyridinium crosslinks. Collagen per construct, represented by hydroxyproline (hypro), pyridinium (PYD) normalized to wet weight (WW) and hydroxyproline, and activity of LOX released to the media from control clamped constructs through 7 weeks of culture normalized to fresh media which did not come in contact with constructs. Significant differences compared to *0 week and +bracket group (p<0.05).

All constructs had significant increases in mature trivalent PYD crosslinks with time in culture, however treatment with LOXL2 produced further significant increases in PYD crosslinks at 6 weeks compared to control constructs, regardless of whether LOXL2 was applied at 2 or 4 weeks (**Figure 5**). Interestingly, when LOXL2 was applied at 2 weeks, there was no increase in PYD crosslinks compared to control constructs by 4 weeks, however when LOXL2 was applied at 4 weeks there was a significant increase in PYD 2 weeks later at 6 weeks.

To further probe why this temporal response in PYD accumulation may occur, we evaluated the amount of active LOX released to the media throughout culture by cells in control constructs (**Figure 5**). We found that cells release significantly more LOX to the media between the first and third week of culture, suggesting cells are producing more LOX while they are forming aligned collagen fibrils. The amount of LOX in the media then reduces over the next few weeks, but significantly increases in the sixth week again. Since there are significantly high concentrations of active LOX in the media during the second week of culture, this may explain why addition of LOXL2 during the second week of culture does not appear to significantly affect collagen crosslinking by 4 weeks.

### 3.5 Tissue mechanics with temporal LOXL2 application

Mirroring collagen organization and PYD accumulation, all constructs had significant improvements in tensile mechanics with time in culture, with LOXL2 treated constructs having further significant improvements over control constructs by 6 weeks (**Figure 6)**. Specifically, at 6 weeks constructs treated with LOXL2 at 2 and 4 weeks had a 3-6 fold significant increase in modulus of the toe and elastic region compared to controls, a significant 2-3 fold increase in strength at the toe-to-linear transition compared to control, and a significant 3-4 fold decrease in transition and ultimate failure strain (**Figure 6**). Addition of LOXL2 at 2 weeks had no significant effects on tissue mechanics by 4 weeks of culture; however, addition of LOXL2 at 4 weeks did produce significant increases in mechanics 2 weeks later. Further, addition of LOXL2 at 4 weeks resulted in significantly higher toe and elastic moduli at 6 weeks compared to constructs treated with LOXL2 at 2 weeks. Ultimately, constructs treated with LOXL2 at 4 weeks developed tensile moduli of ~26 MPa, matching circumferential direction tensile modulus of immature menisci (13-26 MPa, 0–1 week old bovine) [22,51].

**Figure 6:**
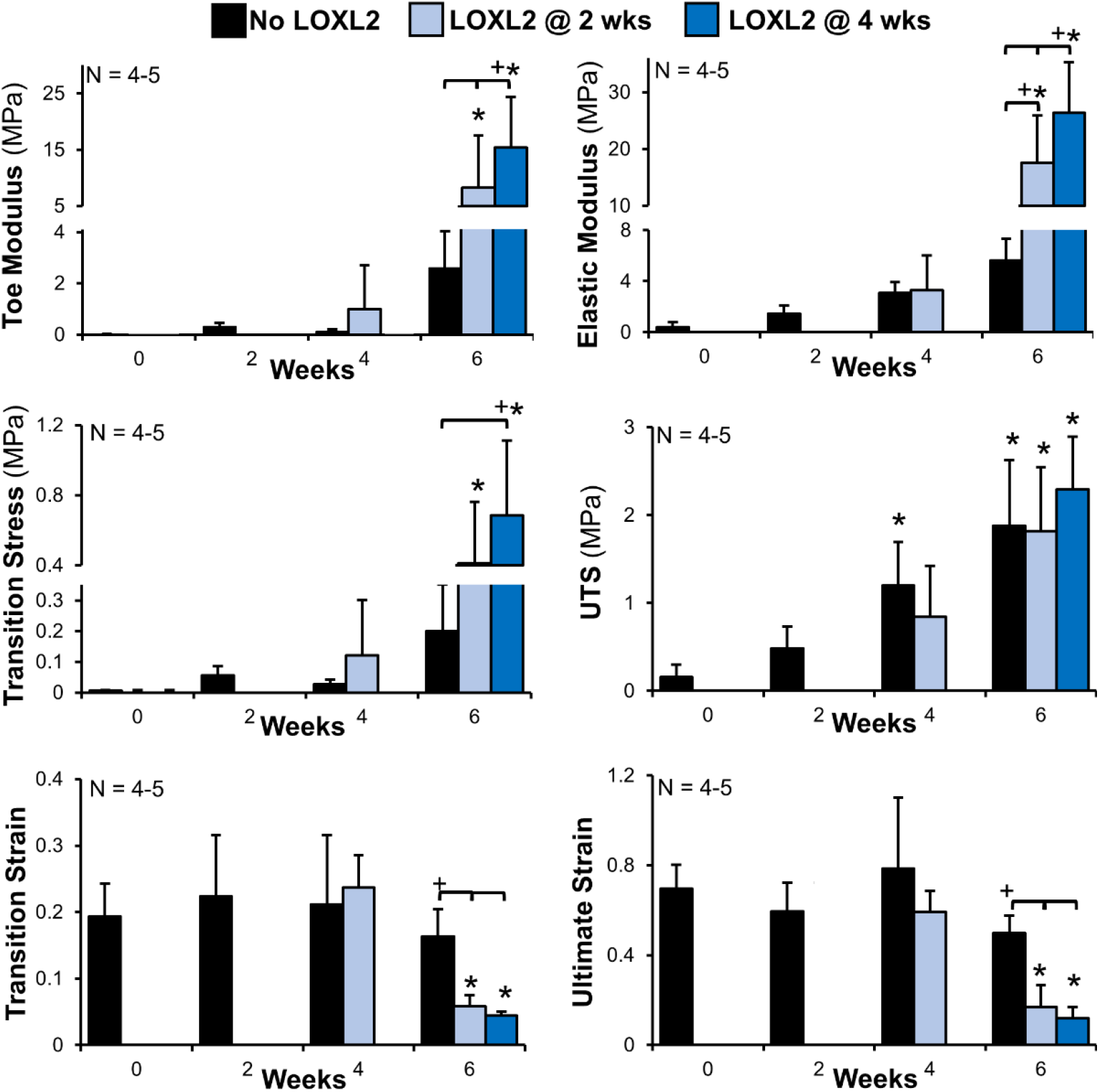
LOXL2 at 2 and 4 weeks of culture produces significant improvements in tensile mechanics. Tensile modulus of the toe region and elastic region, stress and strain at the transition between the toe and elastic region, and ultimate tensile strength (UTS) and ultimate strain of constructs with time in culture. Significant differences compared to *0 week and +bracket group (*p*<0.05).

### 3.6 Application of LOXL2 at 6 weeks

Based off the differences observed with temporal LOXL2 treatment at 2 and 4 weeks, we performed a follow up study where constructs were treated with LOXL2 at 6 weeks once early fascicle formation begins. Previously, we have found that after six weeks cells are often able to rip constructs away from clamps, thus to reduce the risk of early rupture, constructs were only cultured out to 7 weeks. Again, there was no significant difference in gross morphology, collagen organization, or percent weight of control and LOXL2 treated constructs at 7 weeks (**Figure 7 A-C**). Both control and LOXL2 treated constructs had similar organization at the fibril, fiber, and fascicle length-scale, as represented by SEM, confocal, and picrosirius red analysis, respectively (**Figure 7B**). However, LOXL2 treated constructs had a significant decrease in fibril diameter compared to control constructs at 7 weeks, similar to that observed in constructs treated with LOXL2 at 2 and 4 weeks (**Table 1**).

**Figure 7:**
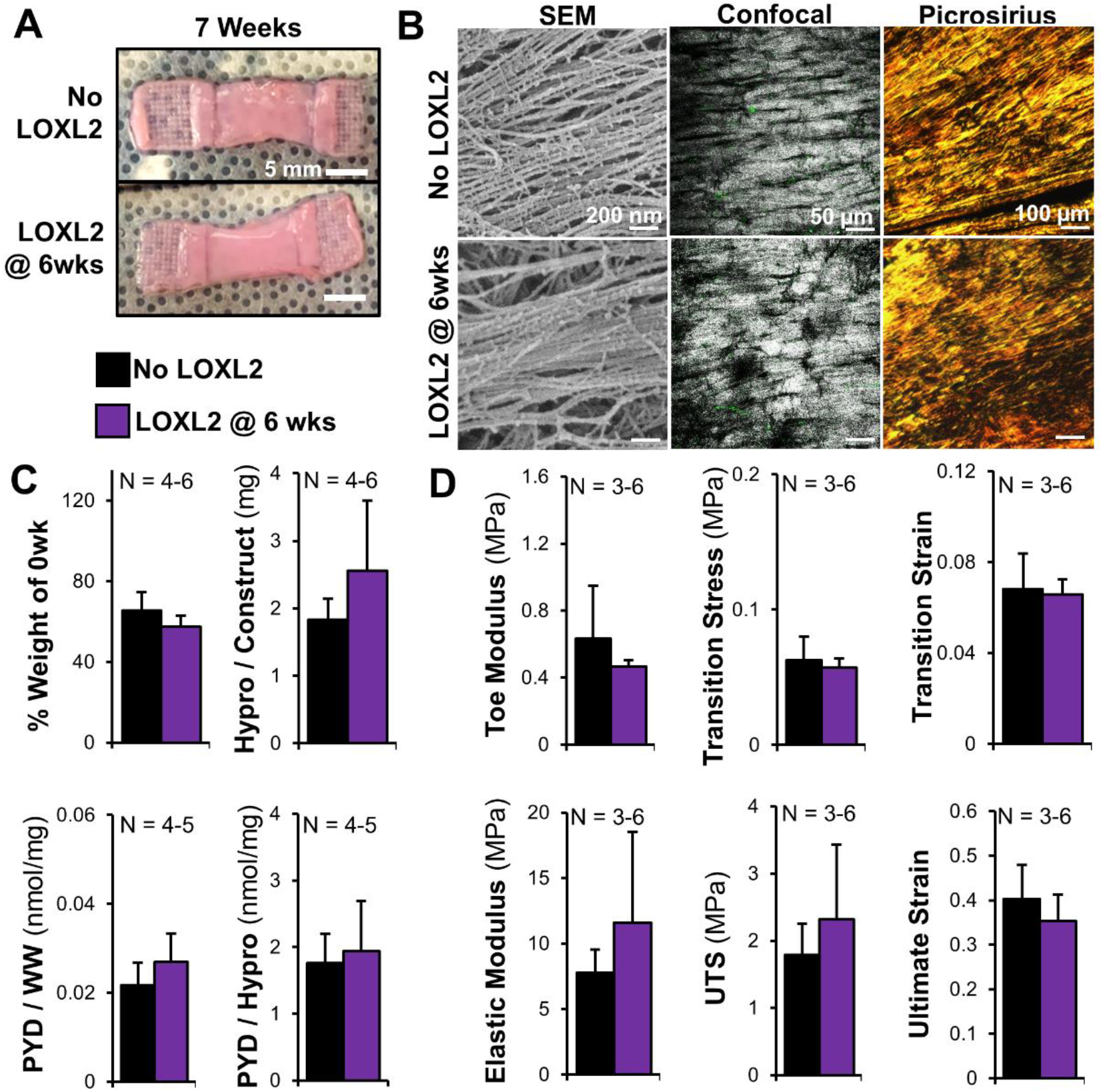
LOXL2 applied at 6 weeks has little effect on organization, composition, and mechanics of constructs by 7 weeks of culture. A) Gross morphology of constructs at 7 weeks. B) Collagen organization of 7 week constructs at the fibril, fiber, and fascicle length scale analyzed via SEM, confocal reflectance, and picrosirius red imaged with polarized light. C) Composition of constructs at 7 weeks, including percent weight of constructs compared to 0 week constructs, hydroxyproline (hypro) per constructs, and pyridinium (PYD) normalized to wet weight (WW) or hypro. D). Mechanical properties of constructs at 7 weeks, including modulus of the toe and elastic region, stress and strain at the transition between the toe and elastic region, and ultimate tensile strength (UTS) and ultimate strain at failure. Significant differences compared to *0 week and +bracket group (*p*<0.05).

Both control and 6 week LOXL2 treated constructs had no significant differences in DNA, GAG, or collagen concentration by 7 weeks (**Supplemental Figure 1C-D and Figure 7C**). Further, both groups had no significant difference in PYD concentration at 7 weeks (**Figure 7C**). Finally, mirroring the minimal differences in collagen organization and PYD accumulation, both control and 6 week LOXL2 treated constructs had no significant differences in tissue tensile mechanics at 7 weeks (**Figure 7D**).

## 4. Discussion

In this study we evaluated the effect of exogenous LOXL2 treatment at different stages of hierarchical collagen fiber formation in an effort to produce functional engineered musculoskeletal replacements and to better understand factors effecting collagen crosslink maturation. We found LOXL2 treatment did not restrict hierarchical fiber formation in our culture device, with constructs still forming aligned collagen fibrils by 2 weeks, larger fibers by 4 weeks, and early fascicles by 6 weeks. However, temporal LOXL2 application did significantly increase mature PYD crosslink accumulation and tissue mechanics. In fact, by 6 weeks constructs treated with LOXL2 once aligned fibrils and fibers developed (2 and 4 weeks, respectively) accumulated ~3 nmol PYD/mg hypro, surpassing values previously reported for immature bovine menisci [51] and reaching ~20-40% of adult bovine and human menisci [38,52]. Further, these LOXL2 treated constructs develop tensile moduli that reached ~26 MPa by 6 weeks, matching immature bovine menisci (13-26 MPa, 0-1 week old bovine) [22,51].

It has been reported that tissue strength is reliant on collagen fiber orientation, density, diameter, and degree of crosslinking [5,23–26]. Previously, we have reported that when culturing cell-seeded high density collagen gels with the boundary conditions in our culture device, cells develop aligned collagen fibrils that match native tissue alignment by 2 weeks and larger fibers that match native fiber diameters by 6 weeks, but mechanical properties plateau at ~1 MPa [19]. In this study our control constructs again develop fibers with similar alignment and diameters as native tissue, and collagen density reached ~200 μg hypro / mg dry weight, matching immature menisci [19]; however, control construct tensile moduli only reached ~5.6 MPa, suggesting the need for further crosslinking. With exogenous LOXL2 supplementation, constructs had a 1.5 fold increase in mature trivalent crosslinks, resulting in a 4-5 fold increase in tensile modulus, demonstrating the importance of trivalent mature crosslinks to tissue mechanics.

Interestingly, we also found that the effect of exogenous LOXL2 application varies depending on the timing of application or degree of hierarchical fiber formation. When LOXL2 was applied at 0 weeks, when collagen was largely unorganized, or at 2 weeks, once aligned fibrils had formed, there was no increase in trivalent crosslinks 2 weeks later; however, when LOXL2 was applied at 4 weeks, once larger fibers had formed, there was a significant increase in trivalent crosslinks 2 weeks later. These differences in kinetics of PYD accumulation could be due to many reasons, including degree of collagen organization, collagen turnover, or competitive inhibition.

The process to make trivalent crosslinks is believed to take weeks to-months *in vivo* [31,33,53], and *in vitro* it has been reported to take 7-30 days for trivalent crosslinks to form in explanted cartilage [50] and 3-4 weeks to form in scaffold free constructs [24,39], both of which largely lack large-scale fiber alignment. However, recent work in chick embryos demonstrated exogenous LOX treatment of calcaneal tendons with highly aligned, developing collagen fibers, produced increased trivalent PYD crosslinking within 3 days [54], suggesting the organization and maturation of collagen plays a significant role in the kinetics of trivalent crosslink formation. It is well established LOX is catalyzed by the local environment and requires collagen fibrils to be aligned in a quarter-stagger array, with crosslinking occurring primarily on the surface of growing fibrils [27–29,31]. Collectively, this suggests the collagen fiber organization and alignment at 4 weeks is more optimally aligned for LOXL2 induced crosslinking, resulting in accelerated accumulation of PYD.

However, in addition to having aligned fibrils, LOX crosslinking is also dependent on collagen molecules having the proper aldehyde precursors or open lysine residues located within the quarter-stagger array of developing fibrils [23,28,30,31]. Tissues throughout the body are well established to have different profiles of LOX crosslinking, with the more highly loaded tissues often accumulating higher concentrations of trivalent crosslinks [28,31,32,35,38]. In fact, positional rat tail tendons are reported to have minimal trivalent crosslinks in comparison to more load-bearing tendons, ligaments, and menisci [28,38,55]. While it is still largely unknown what regulates condensation of immature divalent crosslinks into mature trivalent crosslinks, the location and concentration of hydroxylated lysine residues at the edge of fibrils is thought to play a major role [28,30,31,37,56,57]. Since our high density collagen gels are derived from rat tail tendons, it is possible that the collagen fibrils present at 0 weeks of culture do not possess the proper amino acid sequence or hydroxylated lysine residues to form trivalent crosslinks, accounting for why we do not have increased PYD formation after 2 weeks of culture. Previously, we have found significant release of collagen to the media in the first two weeks of culture [20–22], suggesting cells turn over collagen in the first two weeks, possibly replacing the rat tail collagen with new collagen capable of forming trivalent crosslinks. So PYD accumulation may be accelerated with treatment at 4 weeks not only because the collagen is more organized, but the meniscal fibrochondrocytes may have produced collagen with a more optimal lysine sequence by 4 weeks to better support PYD formation. Future studies should evaluate more long-term time points with and without cells to evaluate whether exogenous LOXL2 treatment at 0 weeks takes longer to form PYD crosslinks *in vitro*, or whether cellular turnover or reorganization of the collagen is needed.

While collagen turnover and reorganization may accelerate PYD accumulation when LOXL2 is applied at 4 weeks, interestingly PYD did not significantly increase when LOXL2 was applied at 6 weeks. A limitation to this study is we were unable to culture these constructs out to 8 weeks to measure whether they accumulated PYD by 2 weeks due to concerns control constructs would rupture; however, within 1 week they did not have significant increases in PYD or tissue mechanics. Recent work has suggested the LOX molecule may be too large to diffuse within the collagen fibril, forcing LOX crosslinking to occur on the outside of growing fibrils [58]. Thus treating constructs with exogenous LOXL2 later in culture, once larger fibers have formed may be less effective at driving trivalent crosslinking or increasing mechanics. Similar work with scaffold free constructs has found early application of LOXL2 during the second week of culture, rather than during the third week of culture, was more beneficial to tissue mechanics [24]. However, the same study found exogenous LOXL2 or hypoxia-induced LOX activity produced increased PYD accumulation and mechanics in explanted 4-8 week old bovine cartilage, tendon, ligament, and meniscus [24], suggesting diffusion of LOX into fibrils may be less of an obstacle in these more immature tissues. Interestingly, this study suggested the time-dependent effect of LOXL2 on mechanics may have more to do with cellular regulation of collagen in the developing constructs [24].

To further probe why this temporal response in PYD accumulation may occur, we evaluated the amount of active LOX released to the media throughout culture by cells in control constructs. We found control constructs had a temporal release in active LOX, with significant increases at 2 and 6 weeks. Thus cells at 2 and 6 weeks may already be making significant amounts of endogenous LOX resulting in limited available lysine residues for exogenous LOXL2 to act on and thus the addition of 150 ng/mL LOXL2 does not make a significant difference. This pattern of LOX expression closely mirrors previously reported results in developing chick calcaneal tendons, where LOX expression, LOX activity, and PYD accumulation increased temporally throughout development, with more dramatic increases at later stages of development [34,36]. Further, this same group has recently demonstrated that treatment with recombinant LOX at different developmental stages results in increased mechanics, with the biggest effect on mechanics occurring with earlier application when endogenous LOX activity is not as high [54]. Specifically, LOX activity and expression is low at Hamburger-Hamilton stage (HH) HH40 and then significantly increases by HH43 [36], similar to what we observe between 4 and 6 weeks of control constructs. When exogenous LOX was applied at HH40 and HH43, there was a larger percent increase in tendon mechanical properties at HH40 compared to application at HH43 [54], again similar to the significant increase in mechanics we observe with LOXL2 applied at 4 weeks vs 6 weeks of culture. This greater increase in mechanics at HH40 was attributed to there being more locations for LOX to crosslink in the developing tendon, prior to the endogenous increase in LOX expression [54].

Aside from temporal differences in response, collectively we found that endogenous LOXL2 increased mechanical properties without significant effects on DNA concentration, collagen concentration, or collagen organization. This matches many previous studies that have found similar results with endogenous application of 150 ng/ml LOXL2 or greater having no effect on cell viability, cell proliferation, collagen concentration, or collagen organization [24,39,41,45,54]. In this study, while addition of LOXL2 at 2 and 4 weeks did not restrict further cellular development of hierarchical fibers later in culture, we did observe a slight increase in distinct fiber and fascicle organization and a significant decrease in fibril diameter for all LOXL2 treated constructs at 6 and 7 weeks. In particular, it appeared that constructs treated with LOXL2 at 4 weeks had more distinct and well organized fibers and fascicles at 6 weeks in comparison to control constructs and constructs treated with LOXL2 at 2 weeks. Previously, when exogenous LOX was applied to developing chick tendons at HH40 and HH43, it was found that HH43 tendons had a significant decrease in collagen dispersion, while HH40 tendons did not [54]. This was attributed to HH43 tendons having denser, highly packed collagen fibers when LOX was applied, allowing for the LOX induced crosslinking to further stabilize the fibers [54]. A similar mechanism may be occurring here when LOXL2 is applied at 4 weeks once larger, denser fibers are formed, in comparison to the smaller fibril organizations observed at 2 weeks. This slight increase in collagen organization may contribute to the fact that constructs treated with LOXL2 at 4 weeks develop stronger mechanics (Toe modulus, elastic modulus, and transition stress) at 6 weeks compared to constructs treated with LOXL2 at 2 weeks, despite both groups having similar concentrations of trivalent crosslinks.

Interestingly, all LOXL2 treated constructs had a significant decrease in fibril diameter at 6 and 7 weeks regardless of timing of LOXL2 treatment. These results correlate with a previous study that found inhibiting LOX with β–amino-proprionitrile (BAPN) during cell driven fibril formation in fibrinogen gels resulted in larger more randomly distributed fibril diameters compared to controls [59]. Collectively, this suggests LOX induced crosslinking helps to tighten or stabilize fibrils in engineered tissues. This may also suggest, that while constructs treated with LOXL2 at 6 weeks did not have increased PYD accumulation, they may have significant increases in divalent crosslinks. As discussed previously, divalent crosslinks are known to form first when LOX catalyzes lysine, and that with time they condense into trivalent crosslinks [28,31,32,50,53]. Divalent crosslinks are reported to form within 1-2 days and often will decrease as trivalent crosslinks form several weeks later [50,53]. However, divalent crosslinks do not have as significant of an effect on tissue mechanics as trivalent crosslinks [28,32]. Thus LOXL2 treatment at 6 weeks may induce divalent crosslinks which do not significantly affect tissue mechanics and by 7 weeks have not had enough time to condense into trivalent PYD crosslinks. Further work should evaluate divalent crosslinks, as well as longer culture durations to determine whether additional time in culture could lead to increased PYD accumulation.

This is the first study to our knowledge to evaluate exogenous LOXL2 treatment at different levels of hierarchical fiber formation, with these constructs holding great promise as engineered replacements for tendons, ligaments, and menisci. Overall, we found 1 week of LOXL2 application at 4 weeks resulted in constructs with native tissue fiber organization and alignment, increased PYD accumulation, and increased tissue mechanics, matching immature bovine menisci. However, further maturation is needed to obtain adult tissue mechanics. A limitation to this study is we only evaluate one concentration of exogenous LOXL2 for our temporal analysis; however, this concentration of 150 ng/ml has been used in many previous studies and shown repeatability to be an ideal concentration for inducing PYD crosslinks in native and engineered tissues [24,40,41,45,46]. We also only dosed constructs with LOXL2 for 1 week at 2 and 4 weeks, and more long-term application or repeated application may be beneficial to tissue maturation. Interestingly, in the initial LOXL2 concentration study we dosed unorganized constructs with LOXL2 for 2 weeks and did not measure an increase in PYD crosslinks, but at 4 weeks of culture in our clamping device, 1 week of LOXL2 treatment produced increased PYD accumulation, suggesting degree of collagen organization or maturation may be more important than dosage duration. Despite these limitations in LOXL2 dosage, constructs require further organization at both the fibril and fascicle length-scale, and increased accumulation of PYD to match adult tissue maturation and mechanics. Our system is a bottom-up approach which mirrors development [19,20]. During postnatal development, as mechanical loading increases, hierarchical organization continues to develop, as does PYD accumulation and mechanical properties [3,26,36,60–62]. Thus, we hypothesize addition of mechanical load to our system will further mature the tissue, resulting in enhanced fiber formation, LOX expression, crosslinking, and ultimately increased mechanics.

### 4.1 Conclusions

Despite the need for further maturation to serve as functional replacements, this study provides new insight into factors that affect mature trivalent crosslink formation. Here we demonstrate LOXL2 can be used to significantly improve the mechanics of tissue engineered constructs, but timing of application is important and will most likely depend on degree of collagen organization or maturation. Currently there is limited understanding of how collagen crosslinking is regulated [30,37], and this system is a promising platform to further investigate cellular regulation of LOX crosslinking. Understanding the mechanism that regulates LOX production and activity is needed to ultimately drive regeneration of functional repair or replacements for connective tissues throughout the body [37].

## Acknowledgements

The authors would like to thank Dr. Carol Mayer and Dr. Dmitry Pestov for their assistance with this study. The authors acknowledge the use of facilities within the Nanomaterials Characterization Core and the Virginia Commonwealth University Cancer Mouse Models Core Laboratory, supported in part, with funding from NIH-NCI Cancer Center Support Grant P30 CA016059.

## Author Contribution

M.B. and J.L.P. conceived the project and wrote the manuscript. M.B. carried out experiments and analyzed the data. L.T. and M.E.B. assisted with SEM and mechanical analysis. J.L.P. supervised the project and acquired funding. All authors edited the manuscript.

