## Supplemental Figure for "Temporal Application of Lysyl Oxidase during Hierarchical Collagen Fiber Formation Differentially Effects Mechanics in Engineered Tissues"

### Supplemental Figures

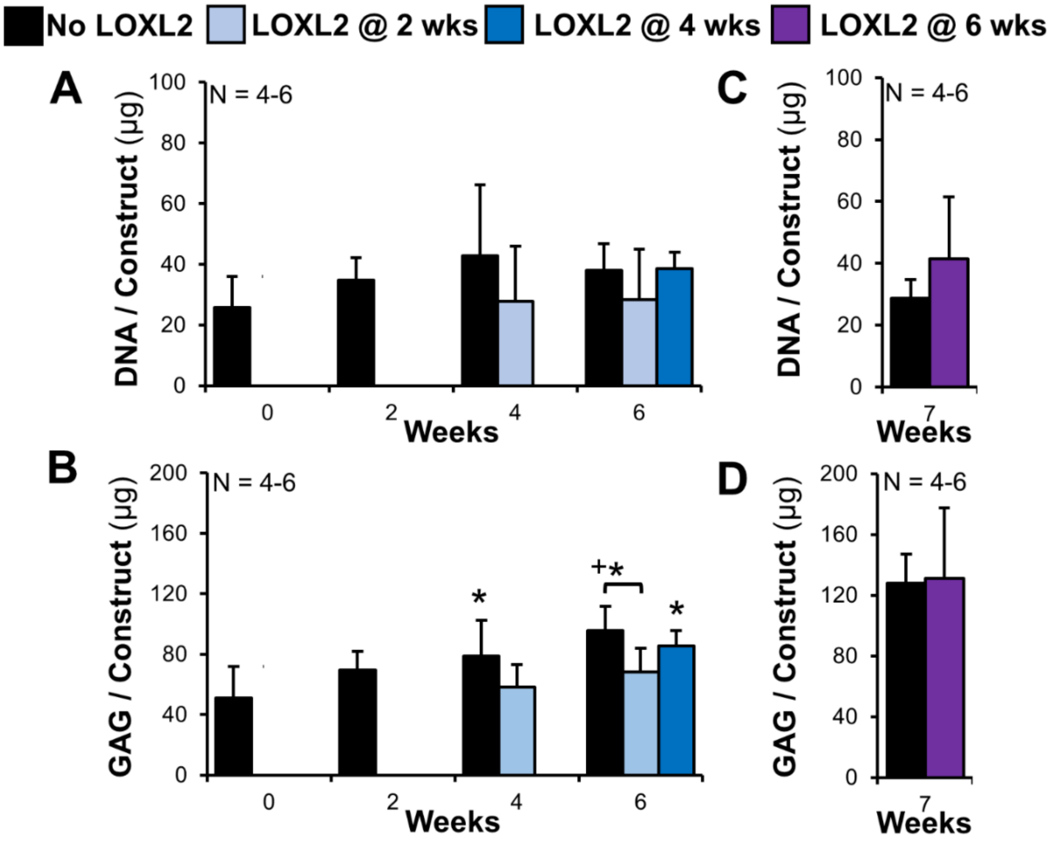


**Supplemental Figure 1:** Temporal application of lysyl oxidase like 2 (LOXL2) has little effect on DNA and GAG accumulation in constructs. DNA and GAG per clamped construct with time in culture when LOXL2 was added at A-B) 2 and 4 weeks of culture and C-D) at 6 weeks of culture. Significant differences compared to *0 week and +bracket group ($p<0.05$). Prior to LOXL2 treatment at 2, 4 or 6 weeks constructs match control (no LOXL2) constructs, thus redundant data prior to LOXL2 supplementation has been removed to allow for better visualization of the data.
